# Artificial selection diversifies birdsong through polygenic changes across sensory and motor systems

**DOI:** 10.64898/2026.10.18.759937

**Authors:** Małgorzata Anna Gazda, William C. Tucker, Paulo Pereira, Pedro Miguel Araújo, Mariel Etchegaray, Larissa B. Patterson, Soraia Barbosa, Raquel O. Vasconcelos, Paulo J. Fonseca, Joel M. Alves, Sandra Afonso, Marisa Ferreira-Marques, Margarida Guimarães-Moreira, Hanka Venselaar, Erwin van Wijk, James Schwartz, Ricardo Jorge Lopes, Tracy A. Larson, Miguel Carneiro

## Abstract

Vocal learning, or the ability to acquire and produce contextualized sounds through social interaction, underlies human speech yet has evolved in a few animal lineages, including songbirds. In songbirds, learned song is produced by a dedicated neural circuit and depends on auditory feedback for its acquisition and maintenance. Roughly four centuries of artificial selection on the domestic canary (*Serinus canaria*) have produced song breeds whose vocalizations diverge as much as those of distinct species, offering a rare intraspecific system in which to identify the genetic and neural changes underlying vocal divergence. Here we show that selection for song reshaped the canary song control system. Compared with breeds selected for plumage or body form, song breeds show increased neural progenitor proliferation and breed-specific changes in neuronal recruitment, sensorimotor brain region volume, and auditory sensitivity. Genome-wide analyses reveal a polygenic architecture underlying song divergence, enriched for transcriptional regulation and synaptic transmission genes. Among these, we find a coding variant in *ADGRV1*, the gene mutated in human Usher syndrome type 2C, associated with low-frequency singing and elevated auditory thresholds in canary. Disrupting *ADGRV1* in zebrafish alters hair cell dynamics, linking adgrv1 mutation to a peripheral sensory mechanism. These findings demonstrate that learned vocal behavior can diversify rapidly through changes spanning sensory, neurogenic, and motor systems, with hearing itself as a substrate for selection.

**Significance Statement:** Centuries of human fascination with the canary’s song have, in effect, conducted a massive parallel evolution experiment on the neural basis of vocal learning. By combining population genomics and comparative neuroanatomy across song breeds with functional tests in zebrafish, we identify molecular and cellular changes by which selection rapidly reshaped learned song. Selection acted not only on the brain but also the ear: a variant in *ADGRV1*, a gene linked to human hearing loss, associates with lower-pitched song and poorer hearing in canaries, and mutation in zebrafish alters sensory hair cell dynamics. The domestic canary reveals how quickly selection can remodel the machinery of vocal learning and offers a tractable system for asking how humans came to speak.

## Introduction

Vocal learning, the ability to acquire and produce vocalizations through imitation, enables elaborate communication, including human speech (*1*). Although this capacity spans a continuum (*2*, *3*), complex vocal learning has evolved in only a few mammalian and avian lineages (*4–6*). Despite independent origins, vocal learning involves similar developmental processes across taxa: a sensory phase in which individuals memorize conspecific sounds and a sensorimotor phase in which they refine their vocalizations through auditory feedback and social interaction (*7*). These processes rely on neural circuits that couple auditory perception with motor control (*2*, *3*, *8*) and are associated with forebrain pathways and gene-expression programs that show convergent similarities across vocal-learning taxa (*9*, *10*).

Although the neural basis of vocal learning is increasingly well defined, its molecular underpinnings remain poorly understood. Learned vocal traits depend strongly on species-specific genetic predispositions (*11*, *12*), yet direct genotype-phenotype links remain rare (*13*). A few genes, including *FOXP2* and *NOVA1*, have been implicated in human speech and vocal evolution (*14*, *15*), and comparative genomics has identified additional candidate loci in accelerated regions of vocal learners’ genomes (*16*, *17*). This gap partly arises because genetic differences in vocal learning are generally found between rather than within species, hindering genetic dissection outside studies of human speech disorders. Systems with heritable intraspecific variation in learned vocal traits therefore offer a direct route to linking genetic variation to behavior.

The domesticated common canary (*Serinus canaria*)—a vocal learner and long-standing model for the behavioral, neural, and hormonal regulation of birdsong—provides such a system (*18*, *19*). Since the 16th century, breeders have selected distinct song types characterized by soft rolling sounds, water-like trills, and bright metallic tones (**Fig. 1A**; **Tables S1**, **S2**). Breed standards, applied both in formal competitions and in breeders’ own selection decisions (**Table S3**), impose sustained directional selection on singing behavior. This process has produced striking acoustic divergence (**Audio S1–S5**). Laboratory crosses and cross-fostering experiments show that breed-specific song traits are heritable, preferentially learned even when alternative tutor songs are available, and determined by a genetic architecture that combines loci of moderate to large effect with additive polygenic effects (*20–23*). The genetic and neural basis underlying these differences remains unknown.

**Figure 1.**
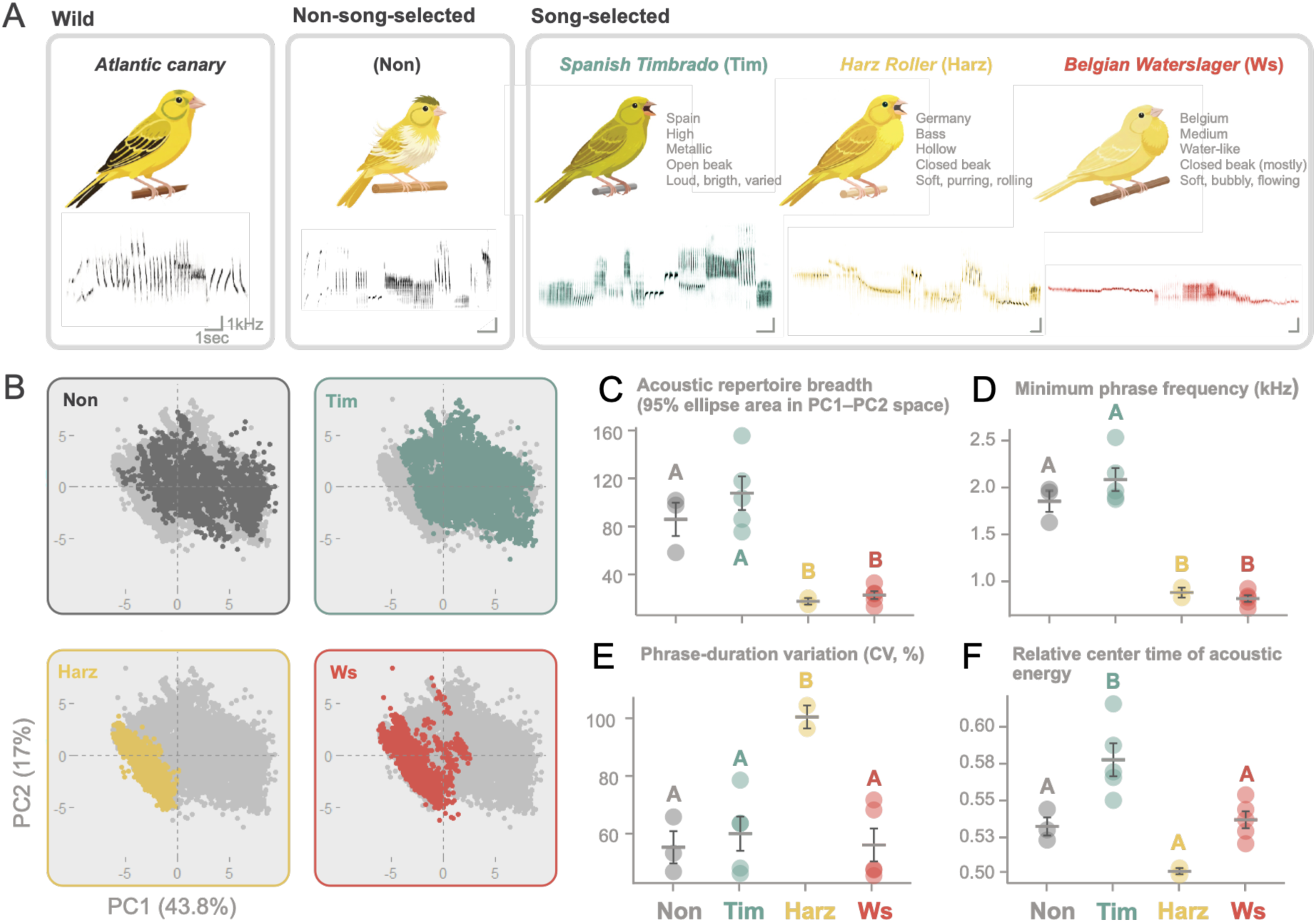
Acoustic differentiation and song structure across canary breeds. (**A**) Representative spectrograms of songs from a wild canary and four domesticated breeds. The wild canary was recorded on Tenerife, Canary Islands (recording: https://xeno-canto.org/791860). Recordings of domesticated breeds were collected in this study. Song-selected breeds include *Harz Roller*, *Spanish Timbrado*, and *Belgian Waterslager*. The *Fife* breed represents the non-song-selected group, which has been selected for morphology and color rather than vocal performance. Labels summarize each breed’s origin, frequency range, timbre, singing style, and overall song character. (**B**) Principal component analysis (PCA) of phrase-level acoustic features. Each panel highlights phrases from one breed in color against phrases from all other breeds (gray), plotted in the shared PC1–PC2 space. Non-song-selected canaries are gray. *Spanish Timbrado*, *Harz Roller*, and *Belgian Watersl*ager are teal, yellow, and red, respectively. (**C**) Acoustic repertoire breadth, measured as the mean area of a 95% probability ellipse in the fixed shared PC1–PC2 space after rarefaction to 100 phrases per bird. (**D**) Minimum phrase frequency (kHz). (**E**) Phrase-duration variation, expressed as coefficient of variation (CV, %). Higher values indicate greater variability in phrase length across a bird’s song. (**F**) Relative center time of acoustic energy (the point within a phrase at which half of the acoustic energy has occurred). (**C–F**) Points represent individual birds; horizontal bars indicate means and error bars indicate SEM. Different letters indicate significant differences among groups based on Tukey tests (*P* < 0.05); groups sharing a letter did not differ significantly. Non-song-selected canaries (Non) are gray, *Spanish Timbrado* (Tim) is teal, *Harz Roller* (Harz) is yellow, and *Belgian Waterslager* (Ws) is red.

Here, we use this long-term “experiment” to investigate mechanisms of song diversification. We combined genetic mapping with quantitative song analyses, neuroanatomical profiling, and functional perturbation to connect genomic variation to divergence in learned vocal behavior. Our findings reveal how strong selection can modify a learned trait, with implications for human speech and other complex behaviors.

## Results

### Artificial selection drives multidimensional divergence in singing behavior

We first investigated acoustic signatures of selection on song. To do so, we compared vocalizations from three breeds selected for singing performance (*Harz Roller*, *Belgian Waterslager*, and *Spanish Timbrado*; hereafter song-selected) with those from canaries bred for color or morphology (hereafter non-song-selected) that retain many ancestral song characteristics of wild canaries (*19*) (**Fig. 1A**). More than 25,000 phrases recorded from 15 birds were used to quantify spectral and temporal features and estimate acoustic repertoire breadth (non-song-selected, *n* = 3; *Harz*, *n* = 2; *Timbrado*, *n* = 5; *Waterslager*, *n* = 5; **Tables S4, S5, Fig. S1**).

Principal component analysis (PCA) on these acoustic features revealed breed-associated differences across acoustic space but without clear separation into breed-specific clusters (**Fig. 1B; Table S6**). This suggests that selection diversified canary song within a common acoustic space, likely constrained by vocal production. To quantify differences in acoustic repertoire breadth among breeds, we measured the space occupied by phrases from each bird in the common PC1–PC2 space. This measure differed among breeds (*F* = 27.79, *P* < 0.0001; **Fig. 1C**). *Harz* and *Waterslager* occupied less acoustic space than *Timbrado* and non-song-selected birds (Tukey tests, all *P* < 0.001), whereas *Timbrado* and non-song-selected birds did not differ (*P* = 0.89). Thus, *Timbrado* exhibited broad acoustic variation, consistent with breed standards emphasizing song diversity, whereas *Harz* and *Waterslager* showed more specialized acoustic distributions (**Tables S1, S2**). We next examined spectral and temporal features underlying song divergence. All 11 frequency measures differed among breeds (ANOVA, all *P* ≤ 0.047; **Fig. 1D**, **Fig. S1**; **Table S5**). For example, *Harz* and *Waterslager* shared a low-frequency spectral profile across the phrase’s lower and upper boundaries and center frequency. This is consistent with breed standards emphasizing low-frequency rolling notes in *Harz* and deep “water-like” sounds in *Waterslager*. By contrast, *Timbrado* differed from all other groups in its low-frequency spectral profile, with higher 5th and 25th frequency percentiles (Tukey tests, all *P* < 0.05; **Fig. S1**; **Table S5**), indicating that *Timbrado* songs contain relatively less energy at the lowest frequencies.

Temporal features also distinguished the breeds. *Harz* showed the greatest variation in phrase duration (ANOVA, *F* = 7.63, *P* = 0.0049; Tukey tests, all *P* < 0.009; **Fig. 1E**), reflecting their characteristic sustained rolling passages. In *Timbrado*, the relative center of acoustic energy (the phrase’s energy midpoint) occurred significantly later than in non-song-selected, *Harz*, and *Waterslager* birds (ANOVA, *F* = 10.61, *P* = 0.0014; Tukey tests, all *P* < 0.022; **Fig. 1F**). Selection in *Timbrado* thus extended to the temporal organization of the phrase. Collectively, these results show that artificial selection has altered the repertoire breadth, spectral properties, and temporal organization of canary song.

### Selection for song shapes auditory sensitivity and song circuit plasticity

To examine the sensory and neural effects of artificial selection on song, we compared auditory sensitivity and song-control circuitry across canary breeds. Using adult male canaries, we first recorded auditory brainstem responses (ABRs), which measure hearing sensitivity along the early auditory pathway. Because altered auditory sensitivity in *Waterslager* is well documented (*23–25*), we used this breed as a benchmark and then examined *Harz* (*n* = 3), *Timbrado* (*n* = 2), and non-song-selected canaries (*n* = 4) using the same approach (**Fig. 2A, Fig. S2**). Similar to the reference *Waterslager* profile, *Harz* birds showed elevated auditory thresholds throughout the tested range (0.5–6 kHz), whereas *Timbrado* and non-song-selected birds had lower, nearly identical profiles. These results extend the *Waterslager* auditory phenotype to *Harz* and highlight lineage-specific changes in auditory sensitivity associated with song selection.

**Figure 2.**
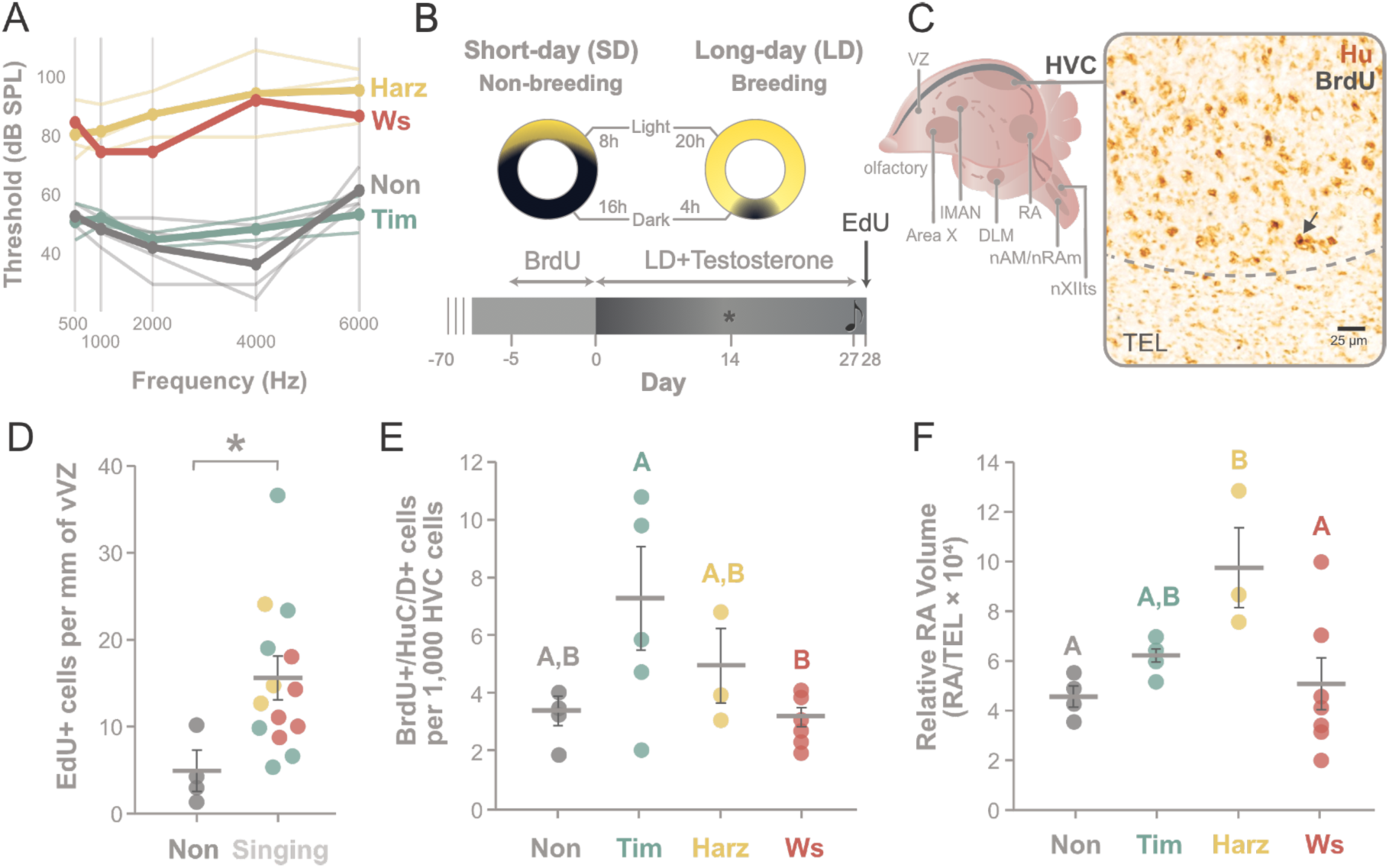
Auditory sensitivity, neural plasticity, and song-circuit morphology in canaries selected for singing performance. (**A**) Auditory brainstem response (ABR) thresholds at five frequencies (0.5, 1, 2, 4, and 6 kHz) in non-song-selected canaries (Non, gray) and the three song-selected breeds: *Spanish Timbrado* (Tim, teal), *Harz Roller* (Harz, yellow), and *Belgian Waterslager* (Ws, red). The threshold was defined as the lowest sound intensity pressure level, expressed in decibels (dB SPL), that elicited a repeatable ABR waveform (**Fig. S2**). Thicker lines show group means. (**B**) Experimental design for inducing breeding condition and associated neural and behavioral plasticity. Birds were implanted with a subcutaneous testosterone pellet and simultaneously transferred to long-day photoperiods (20 h light: 4 h dark) for 28 days. Immediately prior to the onset of breeding condition, birds were injected with the thymidine analog BrdU to label proliferating neural progenitor cells and track their surviving progeny over the following 28–33 days. Song recording began 14 days into breeding condition and continued through the remainder of the experiment (star). Two hours before tissue harvesting, birds were injected with a second thymidine analog EdU to label neural progenitor cells actively proliferating in the ventricular zone at the time of collection. (**C**) Schematic of the songbird brain showing nuclei of the anterior forebrain and descending motor pathways. Regions analyzed in this study include HVC, the adjacent ventricular zone (VZ), RA, and the telencephalon (TEL). Area X, LMAN, DLM, nXIIts, nAM, and nRAm are also shown. On the right, a representative image of brain tissue containing HVC, labeled for the neuronal lineage antigen HuC/D (brown) and the thymidine analog BrdU (black). The arrow indicates a BrdU-positive cell. The full tissue section is shown in **Fig. S3**. (**D**) EdU-positive cells per millimeter of vVZ in non-song-selected canaries and pooled song-selected breeds. Asterisk indicates a significant Welch’s *t*-test comparison. (**E**) BrdU/HuC/D double-positive cells (adult-born neurons) per 1,000 cells in HVC that persisted through breeding condition. (**F**) Relative RA volume, calculated as the ratio of RA to telencephalon volume to control for total brain mass differences. (**D–F)** Points represent individual birds; horizontal bars and error bars indicate means and SEM, respectively. In E and F, different letters indicate significant differences among groups based on Tukey tests (*P* < 0.05); groups sharing a letter did not differ significantly. Non-song-selected canaries (Non) are gray, *Spanish Timbrado* (Tim) is teal, *Harz Roller* (Harz) is yellow, and *Belgian Waterslager* (Ws) is red.

We next characterized neurogenesis, cellular composition, and morphology in HVC and the descending motor pathway of the song-control circuit. This pathway includes projections from HVC to the robust nucleus of the arcopallium (RA) that support the motor control of song production (*26*). Because canaries exhibit seasonal plasticity in song-control nuclei (*26*), we standardized adult male breeding condition using photoperiod manipulation and exogenous testosterone (**Fig. 2B**). We quantified neural progenitor cell (NPC) proliferation in the ventral ventricular zone (vVZ) adjacent to HVC using 5-ethynyl-2′-deoxyuridine (EdU) labeling and recruitment of newly generated HVC neurons using 5-bromo-2′-deoxyuridine (BrdU) and HuC/D double labeling. The vVZ contains neural stem and progenitor cells that supply adult-born neurons to HVC. Cellular composition in HVC was assessed by quantifying HuC/D-positive neuronal and GFAP/vimentin-positive glial populations. We also measured HVC, RA, and telencephalon volumes. These measurements were obtained from 22 birds (non-song-selected, *n* = 4; *Harz*, *n* = 3; *Timbrado*, *n* = 7; *Waterslager*, *n* = 8; **Fig. 2C; Fig. S3**; **Table S7**).

Most cellular and volumetric measures did not differ detectably among breeds (**Table S8**). Three patterns nevertheless emerged. Song-selected breeds collectively exhibited elevated NPC proliferation in the vVZ relative to non-song-selected birds (Welch’s *t*-test, *t* = 3.57, *P* = 0.0045; **Fig. 2D**). This increased neurogenic capacity thus appears to be a shared feature of artificial selection for singing performance. Neuronal recruitment and descending motor pathway morphology exhibited breed-specific specializations. For example, the density of newly generated HVC neurons differed among breeds (ANOVA, *F* = 3.42, *P* = 0.045), with *Timbrado* showing the highest mean density and significantly exceeding *Waterslager* (Tukey test, *P* = 0.041; **Fig. 2E**). This variation points to breed-specific differences in adult-born neuron recruitment or survival and may indicate differences in song circuit plasticity. Relative RA volume also differed among breeds (ANOVA, *F* = 4.44, *P* = 0.020), with Harz showing the largest mean RA to telencephalon ratio, significantly exceeding non-song-selected and *Waterslager* birds (Tukey tests, *P* = 0.024 and *P* = 0.023, respectively; **Fig. 2F**). Selection for song has thus acted on auditory sensitivity, neural plasticity, and motor circuit morphology.

### Genetic architecture of song divergence

To investigate the genetic basis of these differences, we sequenced 75 birds representing the three song-selected breeds, ten non-song-selected breeds/varieties, and wild canaries sampled within their native range in the Canary Islands (**Table S9**). We hypothesized that sustained selection for vocal performance left genomic signatures at loci contributing to song-related traits. To identify candidate regions under selection, we conducted genome-wide scans using DCMS to integrate complementary signals of genetic diversity, population differentiation, and haplotype structure (**Methods**).

We identified 61, 68, and 72 candidate intervals in *Harz*, *Waterslager*, and *Timbrado*, respectively, distributed across multiple chromosomes (**Fig. 3A; Fig. S4; Table S10)**. This genome-wide distribution supports a polygenic architecture of song divergence, consistent with previous crosses (*20*, *22*, *23*). Multiple candidate intervals were intergenic (*Harz* = 23.0%; *Waterslager* = 10.3%; *Timbrado* = 22.2%). Among gene-containing intervals, few contained highly differentiated protein-coding mutations (ΔAF ≥ 0.7); therefore, most intervals lacked either coding sequence or strongly differentiated coding variants (*Harz* = 82.0%, *Waterslager* = 89.7%, *Timbrado* = 98.6%). These observations point to a regulatory rather than protein-coding basis for most candidate regions.

**Figure 3.**
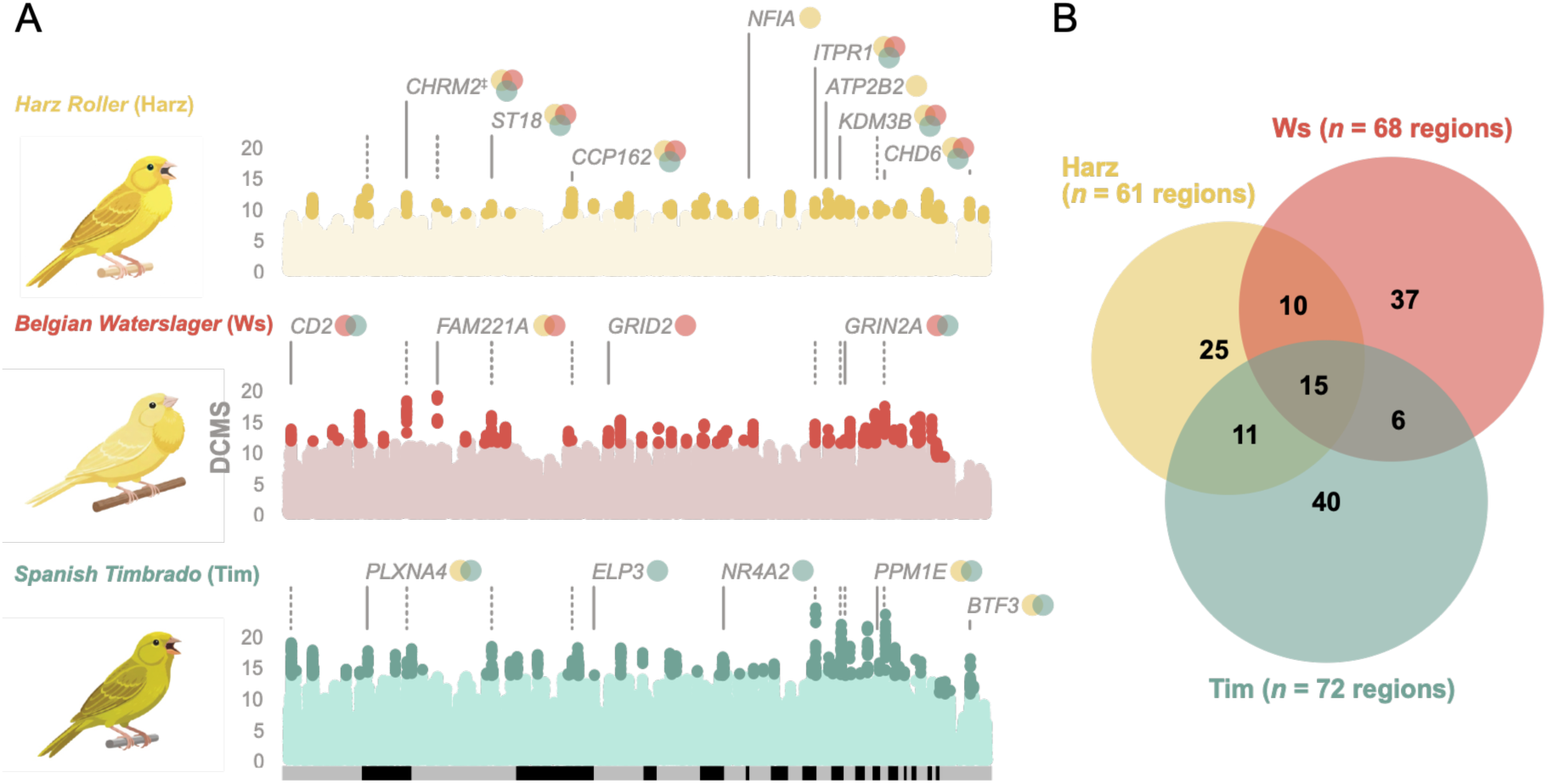
Genomic signatures of selection across canary breeds. (**A**) Manhattan plots of genome-wide DCMS values estimated for each song-selected breed. Each point represents a 50 kb window, with genomic position on the *x*-axis and DCMS value on the *y*-axis (negative values not shown; **Methods**). Darker points indicate windows in the top 0.5% of the DCMS distribution. Solid lines identify annotated genes within candidate regions, with small Venn diagrams beside gene names indicating which breeds share the corresponding candidate region. Dark dashed lines connect the positions of annotated genes shared among breeds. ‡ denotes genes immediately adjacent to, but outside, a candidate region. (**B**) Venn diagram showing the numbers of shared and breed-specific candidate regions across the three song-selected canary breeds.

Next, we quantified the overlap among breed-specific candidate regions (**Fig. 3B)**. Forty-three regions overlapped in at least two breeds and 15 across all three breeds. This level of sharing is highly unlikely under random expectation (all pairwise and three-breed comparisons, *P* < 0.0001). This overlap could reflect: (i) convergent selection, where independent selection acts on the same loci in different breeds; (ii) introgression, driven by secondary gene flow between breeds; or (iii) shared ancestry, where selected variants originated prior to breed divergence.

One candidate region shared across all three breeds contained only *ST18*, a transcription factor involved in neuronal differentiation (*27*). Other shared regions were located near *CHRM2*, which regulates neuronal excitability and is expressed in song-control nuclei (*28*), and *ITPR1*, which is required for synaptic plasticity and motor coordination (*29*). Shared single-gene intervals also contained the chromatin regulators *KDM3B* and *CHD6*. Several candidate regions shared between two song-selected breeds also contained a single annotated gene. For example, a *Waterslager*-*Timbrado* interval contained *GRIN2A*, a gene involved in glutamatergic synaptic plasticity (*30*). *GRIN2A* and other candidates, including *GABRD* and *CHRNB2*, participate in chemical synaptic transmission, the most significantly enriched biological process among genes in candidate intervals (**Table S11**). Two *Harz-Timbrado* regions contained *PLXNA4*, an axon-guidance mediator expressed in vocal-control nuclei (*31*), and *PPM1E*, a regulator of neuronal development that harbors a high-frequency stop-gain variant in *Harz* (*32*).

A substantial fraction of candidate regions was nevertheless unique to each breed (*Waterslager* 54.4%; *Harz* 41.0%; *Timbrado* 55.6%; **Fig. 3B**). Song diversification thus followed partly distinct genetic trajectories across breeds. As with the shared regions, several breed-specific intervals also narrowed to a single annotated gene. *Harz* candidates included *NFIA*, a transcription factor that regulates neural progenitor differentiation and cooperates with *FOXP2* to promote neuronal maturation (*33*), and *ATP2B2*, a calcium pump essential for hair-cell function (*34*). A *Waterslager*-specific interval encompassed *GRID2*, a glutamate receptor required for synaptic plasticity, speech, and motor coordination (*35*). *Timbrado* candidates included *NR4A2*, a regulator of dopaminergic neuron development and motor circuitry (*36*), and *ELP3*, a putative *FOXP2* target in the songbird brain (*37*). These scans thus implicate a functionally diverse set of genes in song divergence, including genes previously implicated in neural plasticity and vocal learning as well as novel candidate regulators of these processes.

### Z-linked mutation associated with low-frequency singing

To identify variants associated with a specific acoustic phenotype, we took advantage of the shared low-frequency singing of *Waterslager* and *Harz*. Previous laboratory crosses suggested that Z-linked factors influence the proportion of low-pitched song elements (*22*), leading to the prediction that at least part of the association would map to the Z chromosome. We therefore performed a genome-wide association analysis comparing the genomes of *Waterslager* and *Harz* (*n* = 25) with *Timbrado*, multiple non-song-selected breeds, and wild canaries (*n* = 50). This design targets genetic factors shared by low-frequency breeds rather than breed-specific alleles.

We identified 235 variants significantly associated with low-frequency singing after Bonferroni correction (**Fig. 4A**). Consistent with the above prediction, 233 (99%) mapped to two regions on the Z chromosome. The first comprised 78 variants spanning ∼155 kb (chrZ:18,285,404–18,440,770 bp) in a noncoding interval between *NUDT12* and *EFNA5*. Whereas *NUDT12* has no clear link to vocal traits, *EFNA5* encodes ephrin-A5, a ligand involved in axon guidance and neural circuit formation (*38*). We found no structural variants strongly differentiated between low-frequency breeds and controls in this interval, so the association may reflect one or more small-scale regulatory sequence variants.

**Figure 4.**
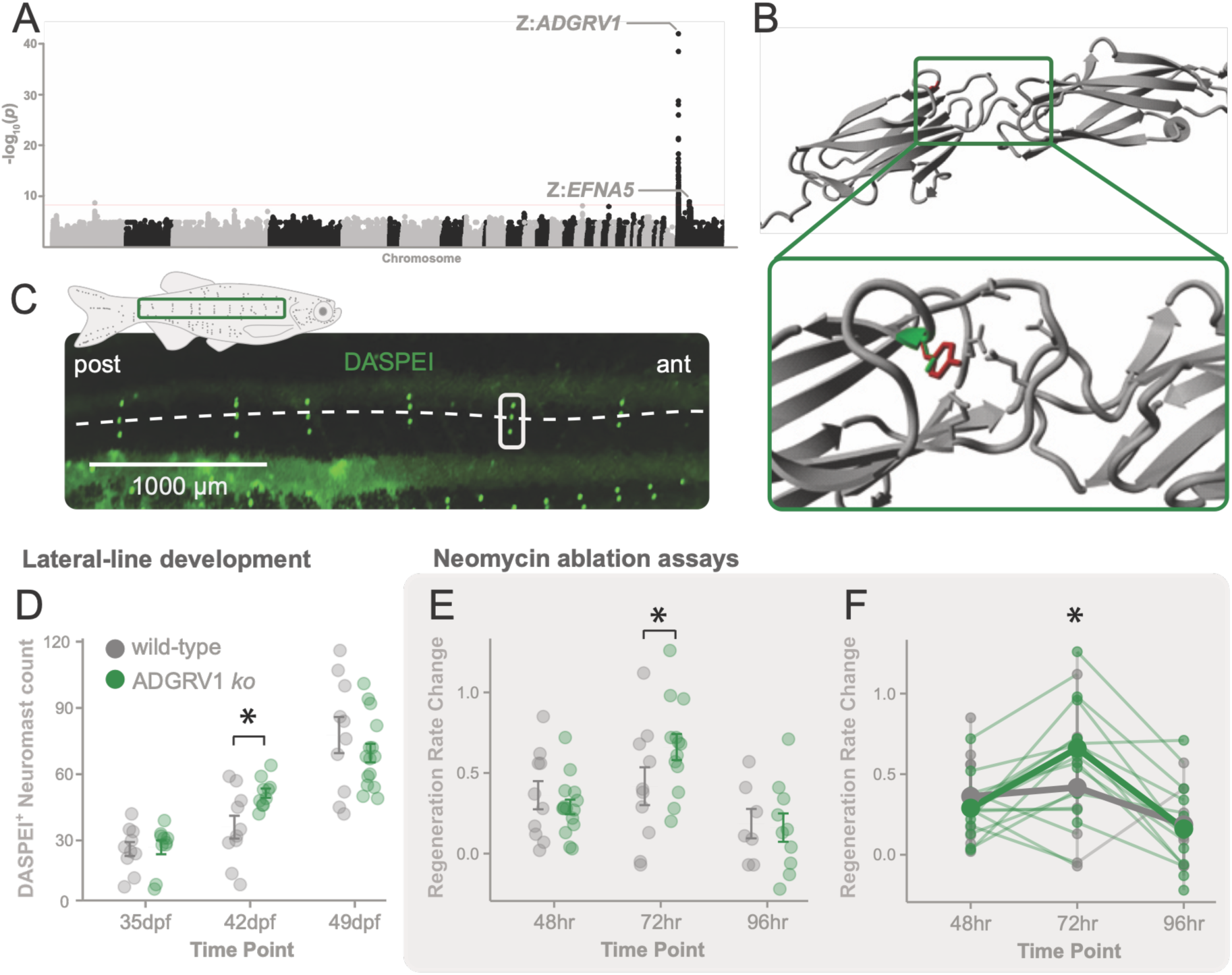
*ADGRV1* is linked to low-frequency singing and altered neuromast dynamics. (**A**) Genome-wide association study (GWAS) results comparing low-frequency singing breeds (*Waterslager* and *Harz*; *n* = 25) to controls (*n* = 50). Each dot represents a tested variant, plotted by chromosomal position (x-axis) and −log10 (*P*) (y-axis). The horizontal line marks the Bonferroni significance threshold. Most significant associations cluster on the Z chromosome at two loci, labeled Z:*EFNA5* and Z:*ADGRV1*. (**B**) Structural context of the D2639Y substitution in ADGRV1. Prediction models of the human ADGRV1 protein show the impact of the candidate canary variant. Wild-type D2639 (green) and mutant Y2639 (red) are located at the interface between two Calxβ domains, in a region predicted to coordinate calcium ions. The side chains of predicted calcium-binding residues are shown in gray. Substitution of tyrosine for aspartate removes a conserved negatively charged residue and may disrupt calcium binding and local domain interactions. (**C**) Representative DASPEI staining (green) of the posterior lateral line in zebrafish, labeling mechanosensory hair cells within neuromast rosettes. The schematic (top left) illustrates the position of the imaged region along the trunk. The dashed line marks the lateral line midline; the white box highlights an example neuromast rosette. Posterior (post) is to the left and anterior (ant) to the right. Scale bar, 1,000 μm. (**D**) Number of DASPEI-positive neuromasts containing hair cells in the posterior lateral line of *adgrv1* CRISPR-mutagenized zebrafish (green circles) and wild-type sibling controls (gray circles) at 35, 42, and 49 days post-fertilization (dpf). Each point represents one fish. Horizontal bars and error bars indicate means and SEM, respectively. Asterisk (*) denotes significant post-hoc pairwise comparisons. (**E–F**) Neuromast regeneration following neomycin-induced hair-cell ablation in *adgrv1* mutants (green) and wild-type controls (gray). Vertical error bars show group means ± SEM. Asterisk (*) denotes significant comparisons. (***E***) Proportion of neuromasts regenerated relative to pre-ablation baseline, measured at 48, 72, and 96 hours post-ablation. Each point represents one fish. (**F**) Individual-level trajectories of neuromast regeneration proportion across the same post-ablation time course. Each line represents a single fish tracked longitudinally.

The second region showed a stronger association (lead variant, *P* = 1.3 × 10^-42^). Spanning ∼460 kb (chrZ:1,183,510–1,642,281 bp), this locus contained 155 significant variants and overlapped five protein-coding genes (*CETN3*, *MBLAC2*, *POLR3G*, *LYSMD3*, and *ADGRV1*). We found no large structural or loss-of-function variants. We did identify two strongly differentiated missense variants in *ADGRV1* between low-frequency breeds and controls (ΔAF = 1.00 at Z:1,388,587; ΔAF = 0.97 at Z:1,435,844). The former was the only one of 96,849 coding variants genome-wide with fixed alternative alleles between groups, and was the lead association signal at this locus. The allele homozygous in *Harz* and *Waterslager* birds was absent from other domestic breeds and wild canaries, suggesting that it arose during domestication. Its presence in both breeds is likely explained by introgression or shared ancestry. The mutation causes a D2639Y substitution (aspartate to tyrosine) at a residue conserved across 619 vertebrate species, underscoring its likely functional importance. Structural modeling places D2639 at the interface between adjacent Calxβ domains, where substitution with a bulkier, neutral tyrosine could disrupt predicted calcium binding and local domain interactions (**Fig. 4B**).

Mutations in *ADGRV1* cause Usher syndrome, a leading cause of combined deafness and blindness in humans (*39*). The elevated hearing thresholds in both low-frequency-singing breeds therefore raise the possibility that D2639Y contributes to their song phenotype by altering auditory function. In *Waterslager*, hair-cell abnormalities have also been reported (*40*). To test the functional consequences of *ADGRV1* disruption in a mechanosensory context, we generated CRISPR/Cas9-induced *adgrv1* mutants in zebrafish and examined the lateral line (**Fig. 4C**). Lateral line surface neuromasts contain hair cells analogous to those of the avian and mammalian inner ear and provide an experimentally accessible system for tracking hair-cell dynamics over time (*41*). We quantified neuromast numbers at seven-day intervals following fertilization (days post-fertilization; dpf) and found significantly increased numbers in mutants following larval-to-adult remodeling at 42 dpf (ANOVA, *F* = 54.17, *P* < 0.0001; post-hoc *P* = 0.002; **Fig. 4D**; **Tables S12**, **S13**), followed by a marginal trend toward fewer neuromasts at 49 dpf (post-hoc pairwise comparison, *P* = 0.073).

We then asked whether *adgrv1* disruption alters regenerative capacity. We ablated mechanosensory hair cells using neomycin, which leaves supporting cells intact, and measured recovery relative to pre-ablation levels. At 72 hours post-ablation, but not at 48 or 96 hours, mutants recovered a greater proportion of neuromasts than wild-type siblings (ANOVA, *F* = 58.63, *P* < 0.0001; post-hoc *P* = 0.036) and showed a higher regeneration rate (ANOVA, *F* = 5.05, *P* = 0.0006; post-hoc *P* = 0.039; **Fig. 4E, F**; **Tables S12**, **S13**). Disruption of *adgrv1* thus alters both neuromast development and regeneration. Together, these findings identify *ADGRV1* as a candidate genetic link between low-frequency singing and auditory sensitivity.

## Discussion

Centuries of artificial selection in canaries have created a unique system for studying birdsong diversification. Previous genetic mapping studies of avian vocal variation have focused on between-species differences in systems with innate vocal development (*43*). By contrast, canary breeds offer heritable variation in learned song within a single species. We demonstrate that rapid song diversification during canary domestication was accompanied by genomic, neuroanatomical, and auditory changes. Rather than arising from individual genetic switches, our findings point to a polygenic basis for song divergence, involving multiple gene functions and biological systems.

Canary song diversification across repertoire breadth, spectral properties, and temporal organization was accompanied by shared and breed-specific modifications of the vocal system. All three song-selected breeds showed elevated neural progenitor proliferation in the ventricular zone that supplies HVC with adult-born neurons, suggesting a shared increase in neurogenic capacity upon which breed-specific specializations were layered. *Timbrado*, selected for a diverse repertoire, showed greater incorporation of new neurons into HVC, consistent with artificial selection acting on a pre-existing relationship between HVC size and vocal complexity documented across species (*44*). *Harz*, characterized by sustained and finely controlled vocal output, had an enlarged RA, the descending motor target of HVC and the avian analog of the human laryngeal motor cortex (*9*), suggesting that control of extended song sequences requires expanded motor capacity. Beyond these neuroanatomical specializations, *Harz* and *Waterslager* exhibited elevated hearing thresholds, consistent with selection for singing behavior modifying auditory function alongside central song-control circuitry.

Shared and breed-specific patterns were also evident in our selection scans, which identified candidate regions across many chromosomes. This inferred polygenic architecture contrasts with avian coloration phenotypes, which often involve a few large-effect loci (*45–47*). The difference may reflect the biological organization of these traits. Coloration can be modified by a few pigment-pathway genes in the integument, whereas song integrates auditory perception, learning, and motor control across many genes and tissues. This diversity also extends to molecular function. Several candidate genes encode transcription factors and chromatin regulators that act at distinct levels of gene regulation. The recurrence of candidate genes associated with expression regulation may reflect selection on developmental programs governing when, where, and how neural cells are generated, specified, and incorporated into song circuits. Chemical synaptic transmission was also the most significantly enriched biological process among candidate genes, indicating that selection targeted both vocal-circuit assembly and the synaptic mechanisms governing their function.

Among these candidates, *ADGRV1* may link peripheral sensory function to the acoustic specialization of low-frequency singing breeds. In mammalian hair cells, ADGRV1 forms ankle links between adjacent stereocilia, and its loss disrupts hair-bundle morphology and mechanotransduction (*48*). In *Waterslager* canaries, elevated hearing thresholds are not congenital, but emerge over the first weeks post-hatch, tracking a progressive decline in hair cell number complete by the onset of song learning (*49*). Our zebrafish experiments also showed that *adgrv1* disruption alters neuromast development and regeneration, with effects emerging during larval-to-adult remodeling. These altered neuromast dynamics are consistent with the concurrent hair-cell death and ongoing regeneration reported in *Waterslager* canaries (*25*, *42*). Together with the established involvement in human Usher syndrome, our zebrafish results support a broader role for *ADGRV1* in mechanosensory hair-cell development, maintenance, and regeneration in vertebrates. However, they do not establish how D2639Y specifically affects the canary ear. We hypothesize that early increases in progenitor proliferation could accelerate hair-cell turnover and prematurely deplete progenitor pools required to maintain auditory sensitivity, contributing to the elevated ABR thresholds in *Harz* and *Waterslager*.

Although birdsong and human speech are not directly comparable traits, our findings offer insight into how learned vocal behaviors can respond to selection. At the molecular level, the scarcity of strongly differentiated coding mutations and the recurrence of candidate genes encoding transcription factors and chromatin regulators suggest an important role for regulatory change in the diversification of learned vocalizations (*9*, *17*). The noncoding region near *EFNA5* provides a striking example, linking low-frequency singing in canaries with a locus known for accelerated noncoding evolution in humans and hummingbirds (*16*). This convergence further raises the possibility that the *EFNA5* locus is a recurrent target of regulatory change in vocal-learning lineages and points to a broader role for axon-guidance modulation in vocal evolution. At the anatomical level, the association of *ADGRV1* with low-frequency singing and altered auditory sensitivity suggests that selection may often drive vocal divergence from the sensory periphery inward, plausibly through the auditory-feedback mechanisms on which vocal learning depends. This interpretation is supported by other observations such as the broader covariation between spectral song properties and auditory performance across songbirds (*50*, *51*). Human speech similarly depends on auditory feedback, where impairment likewise alters vocal production (*52*, *53*). Overall, our findings highlight how learned behavior can be reshaped over short evolutionary timescales through polygenic changes affecting its perception, processing, and production.

## Methods

### Ethics statement

All animal procedures were conducted in accordance with NIH guidelines and were approved by the University of Virginia Institutional Animal Care and Use Committee (Protocols #4170 and #4286) and by the Animal Welfare and Ethics Body of CIBIO/BIOPOLIS (2018_01). Wild canary sampling was carried out under the authorization of a local collecting license granted by the Cabildo de La Palma, Consejería de Medio Ambiente, Transportes y Seguridad y Emergencias. All samples were collected by licensed authors of this study.

### Experimental animals and husbandry

Twenty-two adult male domesticated canaries (*Serinus canaria*) were purchased from the pet trade (non-song-selected, *n* = 4; *Harz*, *n* = 3; *Timbrado*, *n* = 7; *Waterslager*, *n* = 8). The non-song-selected group consisted of *Fife* canaries, a type breed selected primarily for morphology rather than song. The ages of birds ranged from 11 to 25 months at completion of tissue harvesting. Prior to the initiation of experiments, all canaries were housed in indoor group aviaries exposed to short-day photoperiods (SD; 8 hr light, 16 hr dark) for 10 weeks to ensure that the song and reproductive systems returned to nonbreeding states and would be sensitive to the stimulatory effects of long-day photoperiods and testosterone pellet implant.

Zebrafish (*Danio rerio*) of the AB-Zirc line were housed at 28 °C on a light cycle of 14 hr light, 10 hr dark. Larval-stage fish were fed marine rotifers supplemented with Artemac (Reed Mariculture), brine shrimp, and GEMMA Micro (Skretting). Adult-stage fish were fed coarse-grained flake food supplemented with Artemac and brine shrimp. Breeding of zebrafish was performed by natural spawning or *in vitro* fertilization after brief anesthesia using pharmaceutical-grade tricaine methanesulfonate (MS-222, Syndel).

### Hormonal manipulation and cell labeling

To examine the morphology and cytoarchitecture of nuclei in the song circuit, male canaries were implanted subcutaneously above the scapula with a 9-mm Silastic capsule (1.47 mm inner diameter, 1.96 mm outer diameter) filled with crystalline testosterone (T; Sigma). Following pellet implantation, birds were transferred to long-day photoperiods (LD, 20 h light:4 h dark) for 28 days to induce breeding-like growth of the song control nuclei (*54*). To assess the addition of adult-born neurons and astrocytes into the song nucleus HVC, birds received daily intramuscular injections of 5-bromo-2’-deoxyuridine (BrdU, 50 mg/kg, 15 mg/ml in 0.9% NaCl; Sigma) for five consecutive days, beginning five days prior to transition to long-day photoperiod and testosterone treatment (LD photoperiod and T pellet; LDT). To label actively proliferating neural progenitor cells, birds were injected intraperitoneally with a single dose of 5-ethynyl-2′-deoxyuridine (EdU, 40.8 mg/kg, 12.3 mg/ml in 0.9% NaCl, Invitrogen; equimolar to BrdU) two hours before tissue collection (*55*).

### Song recording and acoustic analysis

To analyze song behavior, all birds were housed individually in sound isolation chambers (Industrial Acoustics AC-1) beginning 14 days after LDT onset, the time by which song has achieved the stereotyped structure typical of breeding song (*56–58*). Of the 22 individuals purchased, songs were successfully recorded from 15 birds, with a minimum of 100 phrases per bird required for inclusion (non-song-selected, *n* = 3; *Harz*, *n* = 2; *Timbrado*, *n* = 5; *Waterslager*, *n* = 5). In total, 25,070 phrases were analyzed across these birds (median 1,363 phrases per bird, range 103–4,884). Using in-house Python-based software (developed by J. Burt), vocalizations were continuously recorded throughout the remainder of the experiment. Recording parameters were as follows: amplitude threshold, 70 dB; lower noise frequency, 600 Hz; upper noise frequency, 750 Hz; lower song frequency, 750 Hz; upper song frequency, 7,500 Hz. Using *Raven Pro* (Cornell Lab of Ornithology, v1.6), we quantified the phrase-level acoustic variables listed in **Table S4** for all songs sung on day 27 of LDT. For comparisons of individual acoustic features among breeds, we calculated the mean or coefficient of variation across phrases for each bird, using birds as independent observations. Raw phrase-level measurements are provided in **Table S4** and breed-level summaries in **Table S5**.

Principal component analysis (PCA) was performed in *R* using phrase-level acoustic measurements pooled across all breeds. The acoustic variables were centered and scaled to unit variance before PCA. PCA was performed using the *princomp* function. Separate within-breed PCAs were also performed as supplementary analyses but were not used for the figures or comparisons of acoustic repertoire breadth (**Table S6**). Scores from the combined analysis were plotted in separate panels, highlighting phrases from each breed in color against those from the remaining breeds in grey. Variable contributions to each principal component were calculated by dividing each squared loading by the sum of squared loadings for that component. Contributions to PC1 and PC2 were tabulated and ranked to identify the acoustic features contributing most strongly to each axis (**Table S6**).

To quantify acoustic repertoire breadth while controlling for unequal phrase numbers per individual, 100 phrases per bird were randomly sampled without replacement in each of 500 iterations. This rarefaction procedure standardizes sampling effort and has been used in song-repertoire analyses (*59*). PCA scores were fixed from the combined analysis and were not recalculated during rarefaction. For each iteration, breadth was calculated as the area of a 95% probability ellipse fitted to the sampled phrases in the combined PC1–PC2 space. This covariance-based measure follows the general use of ellipse areas to characterize multivariate niche breadth (*60*). Ellipse area was calculated as 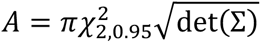, where Σ is the sample covariance matrix of the two PC scores and 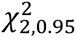 is the 95th percentile of the chi-squared distribution with two degrees of freedom. The mean area across iterations was retained as the bird-level estimate of acoustic repertoire breadth. Bird-level estimates were log-transformed before comparison among breeds using one-way ANOVA.

### Brain extraction and tissue preparation

Six to eight hours after lights-on (Zeitgeber Time 6 to 8), canaries were deeply anesthetized with 4% isoflurane, and brains were rapidly removed. Within two minutes, brains were bisected along the midline. One random hemisphere was frozen on dry ice for cytoarchitectural and immunohistochemical analyses; the contralateral hemisphere was retained for separate analyses not reported in this study. Hemispheres were sectioned in the coronal plane at 40 μm using a cryostat, and sections were thaw-mounted serially. Every third section was Nissl-stained, and the remaining sections were stored at −80 °C until immunolabeling.

### Immunohistochemistry

To visualize BrdU-labeled neurons, we processed brain sections as previously described (dx.doi.org/10.17504/protocols.io.14egn1r8zv5d/v1). Briefly, sections were fixed in 4% paraformaldehyde for 15 min, rinsed in phosphate-buffered saline containing 0.5% DMSO and 0.5% Triton-X (PDTX; pH 7.4), dipped briefly in distilled water, incubated with 2 N HCl at 37 °C for 30 minutes, and rinsed again in PDTX. After blocking in 5% heat-inactivated goat serum, sections were incubated with mouse anti-BrdU (1:350; G3G4 supernatant, Hybridoma Bank) and mouse anti-HuC/D (1:100; antigen 16A11, Invitrogen, A-21271) antibodies. HuC/D (also known as Elav3/4) are neuron-specific antigens expressed in both immature and fully differentiated neurons (*61*). Labeling was visualized with diaminobenzidine (DAB) staining following incubation with biotinylated goat anti-mouse IgG secondary antibodies (1:200; Vector) and amplification with an avidin-biotin peroxidase complex (Vector, VECTASTAIN® Elite ABC-HRP Kit) according to the manufacturer’s instructions. To generate a purple precipitate for BrdU, a DAB-nickel solution was used (0.05% DAB, 0.05% nickel ammonium sulfate, and 0.015% H2O2 in PBS). Neurons were stained with DAB alone to obtain a brown precipitate. Single-and double-labeled cells in each HVC section were counted as described below.

To visualize BrdU-labeled astrocytes, sections were processed as described above, substituting all PDTX rinses with PBS (dx.doi.org/10.17504/protocols.io.261ge1kodv47/v1). Sections were sequentially labeled for BrdU (black) using the antibodies described above and for astrocytes (brown) using rabbit anti-GFAP (1:350, DAKO, Z0334) and mouse anti-vimentin (1:200, Invitrogen, clone V9; 14-9897-82) antibodies. GFAP is expressed in radial glia and mature astrocytes (*62*), whereas vimentin is typically expressed in immature astrocytes (*63*). To label the full astrocyte population in HVC, both antibodies were used in combination and visualized with biotinylated goat anti-rabbit and goat anti-mouse IgG secondary antibodies (both at 1:200; Vector), followed by DAB staining. A distinction between immature and mature astrocytes was not possible with this staining procedure and was therefore not assessed. All astrocytes (brown) and astrocytes double-labeled with BrdU (black and brown) in each HVC section were counted as described below.

To visualize actively proliferative cells, tissue containing the ventricular zone (VZ) was labeled with the Click-iT™ EdU Cell Proliferation Kit for Imaging (Invitrogen, C10337) with minor modifications (dx.doi.org/10.17504/protocols.io.q26g77nb1gwz/v1). The supplied fluorescent azide was replaced with biotin azide (PEG4 carboxamide-6-Azidohexanyl Biotin, Invitrogen, B10184) at a dilution of 1:2000. Otherwise, the Click-iT™ chemistry reaction was performed according to the manufacturer**’**s protocol. Sections were rinsed in PBS and amplified using the avidin–biotin peroxidase complex (Vector, VECTASTAIN® Elite ABC-HRP Kit) according to the manufacturer**’**s instructions. EdU–biotin-azide labeling was visualized using DAB with nickel chromogen as described above. Residual copper from the Click-iT™ reaction can remain bound in tissue following PBS rinses and may react with DAB to produce a greenish-gray precipitate rather than brown. Although this precipitate is quantifiable, it may interfere with additional double-label analyses. If background staining was observed, slides were incubated in 1% EDTA (pH 8.0–8.5) for 10 min following staining to chelate residual copper. All EdU-positive cells were counted along the ventral portion of the ventricular zone (vVZ) as described previously (*64*). For both EdU and BrdU immunohistochemistry, tissue was processed in randomized batches of 30 slides with one slide of fresh-frozen, thaw-mounted EdU or BrdU-labeled gut tissue for positive confirmation of chromogen labeling.

### Morphometric and cytoarchitectural quantification

Volumes of HVC and RA in a randomly chosen hemisphere were determined by tracing the borders of the nuclei identified in Nissl-stained tissue onto paper using a microprojector (Bausch & Lomb). Drawings were scanned, and cross-sectional areas were measured using *ImageJ* (v1.51) (*65*). Total volumes were calculated from traced areas using an *Excel* macro implementing the formula for a truncated cone (*64*). To determine neuronal and astrocyte densities, all DAB-positive cells that fell within an ocular grid (6.25 × 10^-2^ mm^2^ at 400×) positioned in the center of HVC in each tissue section containing HVC were counted. The density of neurons and astrocytes double-labeled for BrdU (DAB-nickel) was quantified using the same procedure. Total numbers of neurons and astrocytes (single-and double-labeled) within HVC were calculated by multiplying cell density for each bird by its unilateral HVC volume. All EdU-positive cells in the ventral VZ were counted from the hillock of the ventral VZ medial to HVC, out to the most lateral extension of the ventral VZ in all sections that exhibited arching of the VZ as previously described (*64*). The total length of the ventral VZ analyzed was measured in *ImageJ*. EdU-positive cells are reported as cells per millimeter (cells/mm) to account for potential differences in ventral VZ size across breeds. Sample sizes varied slightly across specific measures due to tissue staining quality, positive control failure, and subsequent tissue availability (**Tables S7, S8**).

### Auditory brainstem response

The auditory brainstem response (ABR), which reflects activity from the auditory periphery and early central auditory pathways, was recorded from 10 first-year male canaries, including the three song-selected breeds and non-song-selected canaries characterized by wild-type song. Because the auditory phenotype of *Waterslager* has been extensively characterized in previous studies (*23–25*), one *Waterslager* individual was recorded as a reference benchmark to verify agreement with the established elevated-threshold profile. Biological replicates were obtained from three *Harz*, two *Timbrado*, and four non-song-selected canaries for the comparative analysis. Birds for each group were obtained from several local breeders to minimize breeder effects, transported to the laboratory the day before, and housed in cages with food and water *ad libitum*.

During the recording, each bird was sedated with a 10 µL intramuscular injection containing ketamine (50 mg/kg) and diazepam (2 mg/kg) in sterile saline solution (0.9% NaCl) and placed in a prone position on a foam-molded mat that maintained body temperature and position (*23*, *49*, *66*). Recordings were conducted inside a Faraday cage in a dedicated quiet room, with the experimental setup placed on a vibration-isolated table. A speaker was located 30 cm from the bird’s right ear (90° azimuth relative to the bird’s beak; 0° elevation relative to the bird’s right ear). Sound pressure levels were calibrated at the position of the bird’s right ear using a 2238 Mediator Sound Level Meter (Brüel & Kjær) equipped with a Type 4188 ½-inch free-field condenser microphone (Brüel & Kjær), with linear (Z) frequency weighting. Each ABR was recorded using disposable Rochester Horizon™ Subdermal Needle Electrodes (Rochester) with the non-inverting electrode just under the skin at the vertex, the inverting electrode directly behind the right ear canal, and the ground electrode behind the canal of the ear contralateral to stimulation (*49*). Both sound stimulus presentation and ABR recordings were performed using a workstation and software from Tucker-Davis Technologies. Each ABR represented the responses sampled at 20 kHz for 235 ms. The ABR signals were fed from the electrodes into a low-impedance head stage (Medusa4Z amplifier, TDT) connected to a pre-amplifier (RA4PA, Tucker-Davis Technologies) and then routed to a Multi-I/O processor (RZ6, Tucker-Davis Technologies), digitized (16-bit, ± 4 mV) and band-pass filtered (0.03–3 kHz; notch filtered at 50 Hz). Control of sound stimuli and ABR recordings was achieved using *SigGen* and *BioSig* (Tucker-Davis Technologies) software. The acoustic stimuli were 5 ms tone bursts (1 ms rise/fall cos^2^) with a 20 ms inter-stimulus interval, presented at a rate of 4/s. The frequencies 0.5, 1, 2, 4, and 6 kHz were randomly presented at least 1,000 times, half at opposite polarities (180° phase-shifted).

The ABR thresholds for each frequency for each bird were determined by visual inspection of the waveforms by two experienced observers. The threshold was defined as the lowest intensity level at which a repeatable waveform could be identified, based on waveform shape, expected response latency, and, particularly at lower frequencies, the characteristic frequency-doubling response.

### Whole-genome resequencing and read mapping

We obtained whole-genome sequencing data from three domesticated song-selected canary breeds *(Belgian Waterslager*, *Harz Roller*, and *Spanish Timbrado*; *n* = 32 individuals total), ten domesticated color and type canary breeds/varieties (*n* = 33 individuals), and wild canaries from the Canary Islands (*n* = 10 individuals) (**Table S9**). As a source of DNA for genome sequencing, blood was extracted from all birds, both wild and domestic, using brachial venipuncture with sterile needles according to standard collection protocols. The blood samples were preserved in 96% ethanol until DNA extraction. Genomic DNA was isolated using a custom salt-based protocol, followed by RNA removal with RNase A (Roche). Quality control was performed using Nanodrop spectrophotometry for initial purity assessment, and more sensitive fluorometric quantification was conducted with the Qubit dsDNA BR Assay Kit (ThermoFisher Scientific) to ensure high-quality DNA for downstream analysis.

Paired-end sequencing libraries for Illumina sequencing were generated for each of the 75 individuals using the TruSeq DNA PCR-free Library Preparation Kit (Illumina, San Diego, CA). The libraries were sequenced using 2 × 100, 2 × 125, 2 × 150, and 2 × 250 bp reads. The quality of Illumina reads was checked with *FastQC* (https://www.bioinformatics.babraham.ac.uk/projects/fastqc/). Sequencing reads were mapped to the canary reference genome assembly (GCF_022539315.1) with BWA-MEM (v0.7.17-r1188) (*67*) using default settings. To identify duplicate reads, we used the *MarkDuplicates* function from *Picard* (v3.0.0) (http://broadinstitute.github.io/picard). Additionally, *AddOrReplaceReadGroups* from Picard was used to add read group information, which is required for some downstream analyses. Detailed coverage statistics and mapping metrics were calculated using *SAMtools* (v1.11) (*68*) and are provided in **Table S9**.

### SNP and indel variant calling

Variant and genotype calling for SNPs and small indels were carried out using *GATK* (v4.2.6.1) (*69*). Initially, genomic variant call format (gVCF) files were created for each individual employing the *HaplotypeCaller* function, with an additional requirement for a minimum read mapping quality of 30. The resulting gVCF files were then merged into a unified file via the *CombineGVCFs* function, after which the *GenotypeGVCFs* function was used to perform variant calling across all individuals. SNPs and indels were then sorted into distinct files using the function *SelectVariants* and subjected to filtering via the function *VariantFiltration*. The filtration criteria, tailored to each marker type, were applied according to *GATK* documentation:

SNPs:

FisherStrand (FS) > 60.0
QualByDepth (QD) < 2.0
StrandOddsRatio (SOR) > 3.0
RMSMappingQuality (MQ) < 30.0
MappingQualityRankSumTest (MQRankSum) < −12.5
ReadPosRankSumTest (ReadPosRankSum) < −8.0
Variant quality (QUAL) < 30.0
Excess heterozygosity (ExcessHet) > 54.69

Indels:

FisherStrand (FS) > 200.0
QualByDepth (QD) < 2.0
ReadPosRankSumTest (ReadPosRankSum) < −20.0
Variant quality (QUAL) < 30.0
Excess heterozygosity (ExcessHet) > 54.69

SNP and indel variants were then combined into a single VCF file using GATK’s *MergeVcfs* function. Additional filtering steps were applied at both the genotype and variant levels using *VCFtools* (v0.1.16) (*70*). Genotypes with a quality score below 20 (--minGQ 20) and with sequence coverage below 4× or exceeding 48× (twice the average coverage of the individual with the highest coverage) (--minDP 4; --maxDP 48) were classified as missing data. Finally, variants were retained only if they had genotype calls for at least 50% of individuals (--max-missing-count 37). The final dataset for downstream analyses consisted of 7,952,178 variants.

### Variant effect prediction

The protein-coding impact of SNPs and small indel variants was predicted using the Genetic Variant Annotation and Effect Prediction Toolbox (*SnpEff*) (v4.3t) (*71*) and the available annotation of the canary reference genome assembly. For each variant, we calculated the absolute difference in allele frequency (ΔAF) between each song-selected breed and the remaining domestic birds from color and type breeds. Variants with more than one-third of missing genotypes across individuals were excluded. We also recorded for each variant whether alleles were present or absent from wild canaries, identifying those potentially unique to domesticated populations. We concentrated on variants with potential functional consequences, including nonsynonymous mutations, frameshift insertions or deletions, STOP codon gains or losses, and mutations affecting splice sites, as these may have direct impacts on protein structure and function.

### Detection of structural variants

Structural variants were identified and genotyped using the *smoove* (v0.2.8) pipeline (https://brentp.github.io/post/smoove/). This pipeline integrates several tools and performs several read-filtering steps to reduce noise and false positives associated with short-read data. The pipeline begins by calling structural variants for each individual separately, followed by merging the union of detected sites across all individuals. Finally, each individual is genotyped at the shared variant sites. Variants with more than one-third of missing genotypes were discarded. As with the SNP/indel analysis, we calculated for each variant the ΔAF between each song-selected breed and non-song-selected domesticated canaries and retained all variants with ΔAF ≥ 0.70. Promising candidate variants were manually reviewed using IGV (Integrative Genomics Viewer) (v2.16.0) (*72*) for validation, and their locations were cross-referenced with gene annotations provided in the canary reference genome assembly.

### Molecular signatures of artificial selection

Molecular signatures of positive selection across the genomes of each song-selected breed were inferred using three complementary statistics. First, we scanned the genome for locally elevated levels of genetic differentiation by calculating the population branch excess (*PBE*) statistic (*73*). *PBE* is derived from the population branch statistic (*PBS*) (*74*), which contrasts allele frequency divergence between two populations relative to a third, more distantly related population. *PBE* extends this framework by evaluating whether the branch length leading to a focal population at a given locus exceeds expectations based on genome-wide patterns, thereby identifying loci with breed-specific excess divergence. For each analysis, a song-selected breed was designated as the focal population, non-song-selected birds from multiple breeds served as the closely related comparison, and the wild population from the Canary Islands (the species’ native range) was used as the outgroup. Pairwise Weir–Cockerham *F_ST_* values were calculated using *VCFtools* (v0.1.16) in 50 kb sliding windows with 10 kb steps. *F_ST_* values were log-transformed and converted to branch length estimates, and *PBE* (**Fig. S4**) was computed using a custom Python script following Shpak *et al.* (2024) (*75*).

Second, we screened the genome for regions with extended haplotypes at high frequency using the cross-population extended haplotype homozygosity statistic (*XP-EHH*) (*76*). Before running *XP-EHH*, we performed haplotype inference using *Beagle* (v5.1) (*77*) on the filtered set of variants and genotype calls obtained from *GATK* (see above). *XP-EHH* was calculated using *selscan* (v2.0.2) (*78*) by defining each song-selected breed as the target population and both type and color domesticated canaries as the reference population. Given the absence of a canary-specific recombination map, we adopted a recombination rate of 1.5 cM/Mb, based on zebra finch data (*79*). To account for natural variation in haplotype length across the genome, the raw scores at each site with a minor allele frequency higher than 0.05 were normalized using the genome-wide empirical estimates with the function *norm*. Finally, we divided the genome into non-overlapping 50 kb regions and computed the maximum *XP-EHH* score in each window which was then assigned as the entire region’s score (**Fig. S4**).

Lastly, we employed the cross-population number of segregating sites by length statistic (*XP-nSL*) (*80*), which contrasts haplotype lengths and polymorphism between populations, but does not require a recombination map, making it robust to low-recombination regions and better suited for detecting soft sweeps (*80*, *81*). *XP-nSL* was calculated using *selscan* by defining each song-selected breed as the target population and the type and color domesticated canaries as the reference population. We employed the same normalization procedure and window approach as described for *XP-EHH*. In both *XP-EHH* and *XP-nSL*, positive scores correspond to long homozygous haplotypes and putative molecular signatures of positive selection in song-selected canaries versus the other domesticated canaries. Negative values correspond to the opposite haplotype pattern (i.e., indicating selection in the reference population) and were transformed to zero for data visualization (**Fig. S4**).

To combine the information from the three measures of selection signal, while accounting for correlations among them, we applied a De-correlated Composite of Multiple Selection Signals (DCMS) approach (*82*, *83*). In order to calculate *DCMS* we used *PBE*, *XP-EHH* and *XP-nSL*. First, we calculated a p-value for each statistic by sorting their raw, untransformed values in descending order, ranking them, and dividing by the number of observations. Because *XP-EHH* and *XP-nSL* were computed in non-overlapping windows, whereas *PBE* was calculated with a sliding window, each *PBE* value was paired with the closest overlapping *XP-EHH* and *XP-nSL* windows. In parallel, we calculated a correlation matrix across the three statistics by estimating a covariance matrix (*cov* function in *R* using “*pairwise.complete.obs*”) and dividing it by the square root of the outer product of the covariance diagonal with itself. DCMS values were then calculated for each window following Ma et al. (2015). Although implemented in a custom *R* script, the procedure was inspired by the implementation within *MINOTAUR* (*83*). Windows were ranked by DCMS in descending order, and the top 0.5% were designated as candidate regions. All *P*-values, correlation matrices, DCMS, and DCMS significant cutoffs were calculated separately for autosomes and the Z chromosome. For visualization, DCMS with negative values were not shown as they represent regions with a *P*-value close to 1 across all three statistics.

A catalog of regions under positive selection was produced for each breed by merging adjacent candidate DCMS windows along each chromosome, allowing a maximum gap of one window. This procedure was implemented in Python. To measure the overlap between different breeds, a k-way merge of sorted interval lists with overlap coalescing was implemented in Python. Overlaps were then counted and summarized for visualization (**Table S10**).

To evaluate whether the observed number of overlapping regions exceeded the expectations of overlaps obtained by chance, we performed 100,000 permutations. In each iteration, three sets of random regions were generated, matching the number and length distribution of the observed breed-specific regions. Pairwise and three-way overlaps were then recalculated. Empirical *P*-values were obtained as the proportion of permutations yielding an equal or greater number of overlaps than observed.

To prioritize candidates for discussion, we applied three criteria. First, we focused on intervals containing a single annotated protein-coding gene, as such regions provide clearer interpretation by minimizing ambiguity about the likely target of selection. Second, we prioritized genes with established roles in neural development, synaptic function, neuromodulatory signaling, calcium-dependent plasticity, or chromatin and transcriptional regulation in the nervous system. Third, we considered conceptual alignment with our phenotypic data, highlighting genes whose known biological functions could plausibly contribute to differences in auditory sensitivity, neuronal proliferation, circuit plasticity, or motor output underlying song behavior.

### Genome-wide association analysis

Association analyses were conducted using *GEMMA* (v0.98.1) (*84*). We used a linear mixed model and mitigated potential confounding effects arising from population stratification and relatedness among individuals by incorporating in the model: 1) a kinship matrix using the centered relatedness matrix option (-gk 1) in *GEMMA* as a random effect; 2) the first two principal component axes derived from a principal components analysis (PCA) performed using *PLINK* (v1.90b6.26) (*85*) as covariates. To reduce the potential impact of linkage disequilibrium (LD) on kinship matrix estimation and PCA axes, we performed variant pruning in *PLINK* using an *r*^2^ threshold of 0.3 and a window size of 200 kb. Variants with a minor allele frequency below 5% were discarded. Significance thresholds for the association tests were established using a Bonferroni correction adjusted for the total number of tests (0.05/7,430,386 ≈ 6.7 × 10^−9^; corresponding to −log_10_ (*p*) ≈ 8.17).

### ADGRV1 protein modeling and sequence conservation

To assess evolutionary conservation of the candidate mutation, we retrieved ADGRV1 protein sequences from NCBI representing 619 vertebrate species spanning major vertebrate groups, including birds, mammals, reptiles, amphibians, and fish. Sequences were aligned using *FAMSA* (v2.4.1) (*86*) and inspected in *BioEdit* (v7.7.1) (*87*) to evaluate conservation at the mutated residue. To examine the structural impact of the D2639Y substitution (see Results), we generated a model of the ADGRV1 protein using the *AlphaFold server* (https://alphafoldserver.com/) (*88*). The D2639Y mutation was introduced *in silico* using *YASARA* (*89*) to compare the predicted structural consequences of the wild-type and mutant residues.

### CRISPR mutagenesis in zebrafish

CRISPR/Cas9-mediated mutagenesis of *adgrv1* was performed in zebrafish using Alt-R CRISPR-Cas9 reagents (Integrated DNA Technologies). Exon 28 of *adgrv1* was targeted with the crRNA Alt-R 561 (sequence provided in **Table S14**), which was complexed with Alt-R tracrRNA and Alt-R Cas9 Nuclease V3 to form ribonucleoprotein (RNP) complexes (*90*). RNP complexes were microinjected into one-cell stage embryos of the AB-Zirc line. Injected embryos and uninjected siblings were raised to adulthood to generate F0 founders.

To assess mutagenesis efficiency in F1 embryos, pools of four to five embryos were collected and genomic DNA was extracted using a hot-base extraction method. Briefly, embryos were placed in 25 μL of 10% Hank’s solution in PCR tubes, mixed with 25 μL of 2× Base Solution, and heated at 95 °C for 30 minutes. After cooling to room temperature, 50 μL of Neutralization Buffer was added. Genomic DNA was stored at −20 °C or used immediately for PCR (*91*).

The *adgrv1* target site was amplified by PCR in a 25 μL reaction containing 12.5 μL 2× Taq Master Mix (New England Biolabs, M0270L), 1 μL pooled forward and reverse primers (10 μM each), 2 μL genomic DNA, and 9.5 μL nuclease-free water. Thermocycling conditions were: 95 °C for 30 s; 39 cycles of 95 °C for 30 s, 54 °C for 30 s, and 68 °C for 30 s; followed by a final extension at 68 °C for 5 min. Primers flanking a 318 bp region surrounding the Alt-R 561 target site (adgrv1_AltR1-g318f and adgrv1_AltR1-g318r) are listed in **Table S14**. PCR products were purified and subjected to Sanger sequencing. Mutagenesis efficiency and indel spectra were quantified using *TIDE* (Tracking of Indels by Decomposition; https://tide.nki.nl), comparing injected samples with uninjected wild-type sibling controls. F0 mutagenesis efficiency for Alt-R 561 was approximately 71%.

Six F0 founders exhibiting severely degenerated lateral line phenotypes were group-spawned to generate F1 embryos. Genomic DNA was extracted from pools of 4–5 F1 embryos using the hot-base method described above, and the Alt-R 561 target region was amplified and sequenced as described. *TIDE* analysis of pooled F1 embryos revealed segregating mutations, including a predominant 10 bp deletion (∼50% of alleles), a 5 bp deletion (17%), and a 3 bp insertion (20.8%) (**Table S14**). The total editing efficiency in the F1 pools was approximately 92%, indicating successful germline transmission.

### Neuromast characterization in development

To assess lateral line neuromast development, CRISPR-mutagenized fish and sibling-matched controls were stained weekly with the vital mitochondrial dye DASPEI (2-(4-dimethylaminostyryl)-N-ethylpyridinium iodide; Invitrogen) from 7 to 49 days post-fertilization (dpf). At each time point, fish were incubated in 0.005% DASPEI in system water for 15 min at room temperature and rinsed three times in fresh system water to remove excess dye. Stained fish were anesthetized using pharmaceutical-grade tricaine methanesulfonate (MS-222, Syndel) and placed in a glass-bottom imaging dish.

Neuromasts along the full length of the posterior lateral line (trunk and tail) were counted using a Zeiss AxioZoom and AxioObserver microscope equipped with epifluorescence and a Zeiss Axiocam camera controlled by *ZEN Blue* software. The total number of fluorescently labeled neuromasts was quantified by systematic counting along the posterior lateral line. After imaging, fish were recovered in fresh system water and returned to standard group-rearing conditions. The same fish were imaged weekly over the seven-week developmental period to monitor neuromast dynamics over time.

### Neomycin-induced hair cell ablation and regeneration

To assess the recovery of DASPEI-positive neuromasts following neomycin-induced hair-cell ablation, F1 fish and wild-type sibling controls were stained, anesthetized, and imaged at 7 weeks post-fertilization as described above. After baseline neuromast counts were obtained, *adgrv1* mutants and sibling controls were exposed to 500 µM neomycin (Sigma-Aldrich; dissolved in system water) for 1 h to ablate lateral line hair cells (*92*). Staining, anesthesia, imaging, and quantification of DASPEI-positive neuromasts were repeated at 48 h, 72 h, and 96 h following neomycin treatment.

### General statistical analyses

Canary acoustic, morphometric, and cytoarchitectural measures were compared among the four breeds using one-way ANOVAs with breed as the factor, followed by Tukey tests for pairwise comparisons. Morphometric and cytoarchitectural measures were also compared between pooled song-selected and non-song-selected birds using two-sided Welch’s t-tests. Acoustic repertoire areas were log-transformed before analysis. Comparisons of neuromast counts in zebrafish were performed using two-way ANOVA with genotype and time as factors. Post-hoc pairwise comparisons were conducted within each time point to assess differences between mutants and sibling controls. All data are presented as mean ± SEM. Statistical significance was defined as *P* < 0.05. All analyses were performed using *JMP* (v18.0.1; SAS Institute).

## Acknowledgements

We thank numerous breeders who have donated samples. We thank D. Parichy for intellectual guidance and fish husbandry support with CRISPR experiments. This work was supported by: NSF NRT-ROL 2021791 EXPAND Fellowship to WCT; the Portuguese Foundation for Science and Technology (FCT) research contract 2020.01494.CEECIND to PMA; NIH R35 GM150562 to TAL; FCT research contract CEECINST/00014/2018/CP1512/CT0002 to MC; the project PTDC/BIA-EVL/31569/2017 - NORTE-01- 0145-FEDER-30288, co-funded by NORTE2020 through Portugal 2020 and FEDER Funds, and by National Funds through FCT to CE3C (UID/00329/2025, https://doi.org/10.54499/UID/00329/2025).

## Author contributions

Conceptualization: MAG, TAL, MC Methodology: MAG, PP, HV, EW, TAL, MC

Investigation: MAG, WCT, PP, PMA, ME, LBP, SB, RV, PF, JA, SA, MFM, MGM, HV, EW, JS, RJL, TAL, MC

Visualization: MAG, WCT, PP, SB, HV, RJL, TAL Funding acquisition: TAL, MC

Project administration: MAG, TAL, MC Supervision: TAL, MC

Writing – original draft: MAG, TAL, MC

Writing – review & editing: MAG, WCT, PP, PMA, ME, LBP, SB, RV, PF, JA, SA, MFM, MGM, HV, EW, JS, RJL, TAL, MC

