## Supplemental Figures for "Artificial selection diversifies birdsong through polygenic changes across sensory and motor systems"

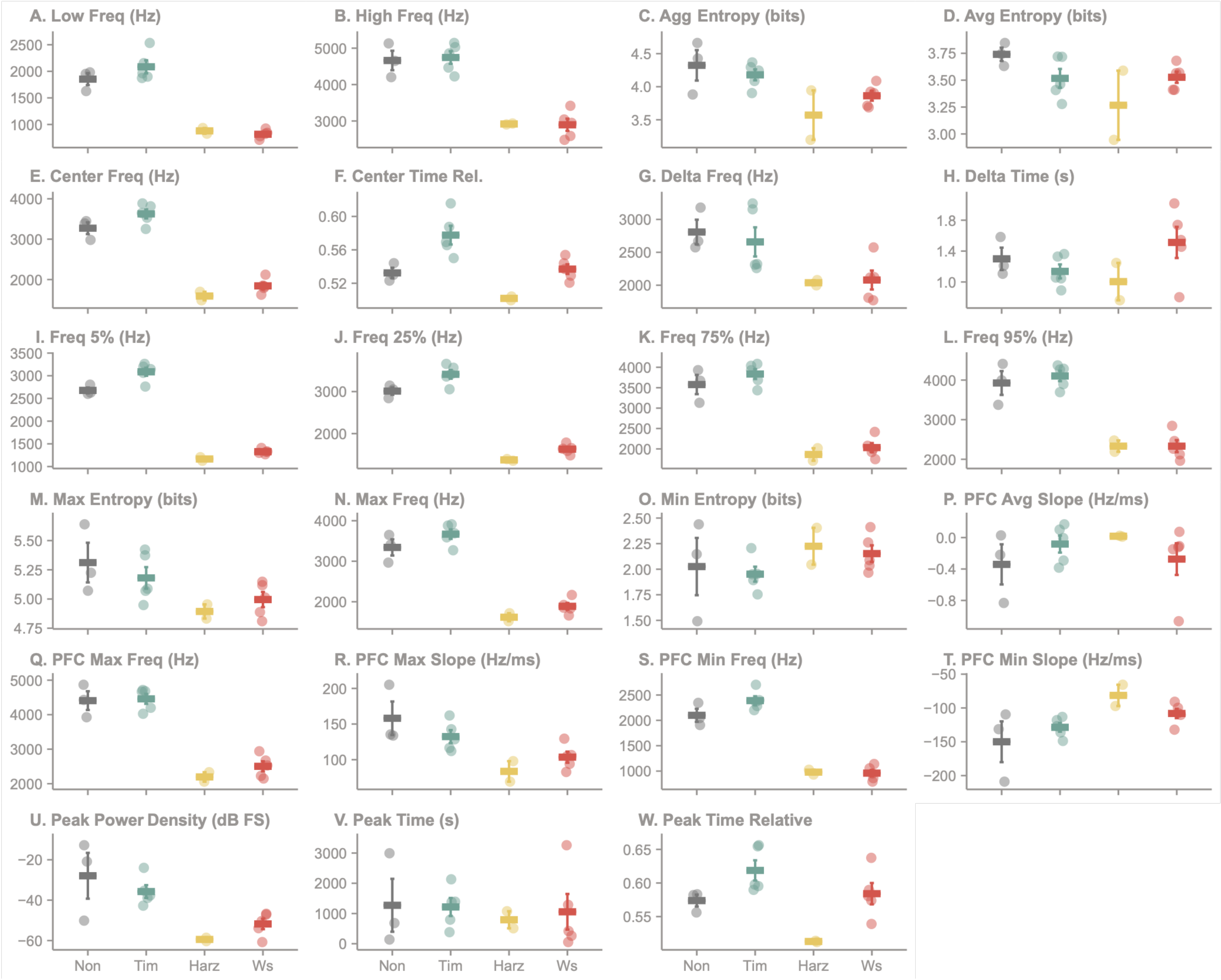


**Figure S1. Phrase-level acoustic features of canary song across breeds.** Each point represents the mean value for one bird across its recorded phrases. Horizontal bars and error bars indicate group means ± SEM. Summary statistics and statistical comparisons are provided in **Table S5**, and raw phrase-level measurements are provided in **Table S4.** Non-song-selected canaries (Non) are gray, *Spanish Timbrado* (Tim) is teal, *Harz Roller* (Harz) is yellow, and *Belgian Waterslager* (Ws) is red.

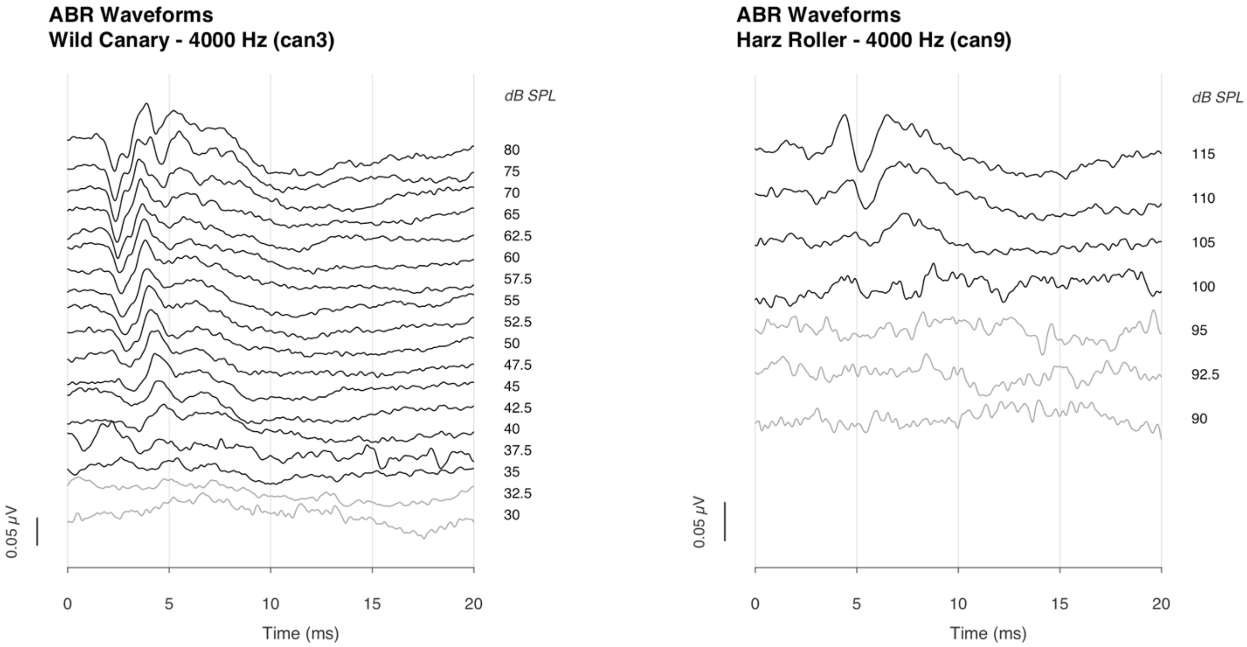


**Figure S2. Representative auditory brainstem response waveforms.** Representative auditory brainstem response (ABR) waveforms recorded in response to a 4 kHz stimulus in a non-song-selected canary (left) and a *Harz Roller* canary (right). Traces show responses across decreasing sound pressure levels (dB SPL), with stimulus intensity indicated to the right of each waveform. Black traces indicate responses with a detectable ABR, whereas gray traces indicate responses below the estimated hearing threshold. The *x*-axis shows time after stimulus onset (ms), and the *y*-axis shows response amplitude (μV).

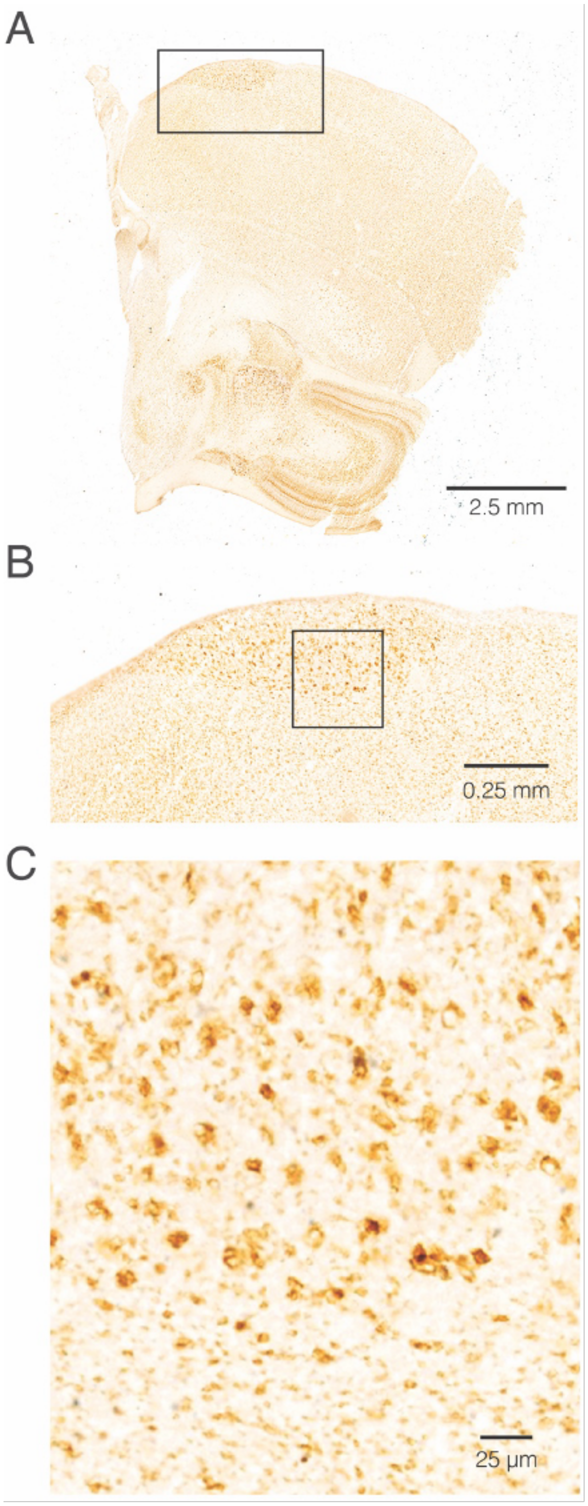


**Figure S3. Representative images of immunohistochemically labeled neural tissue.** The song-control nucleus HVC was labeled with the neuronal lineage antigen HuC/D (brown) and with BrdU, which marks proliferating cells and their recent progeny (black). (**A**) Overview of the brain section containing HVC; the boxed region is enlarged in (B). (**B**) Higher-magnification view of HVC; the boxed region is enlarged in (C). (**C**) Higher-magnification view of BrdU-labeled cells within HVC. The corresponding image without annotations is shown in **Fig. 2C**. Scale bars: (A), 2.5 mm; (B), 0.25 mm; (C), 25 µm.

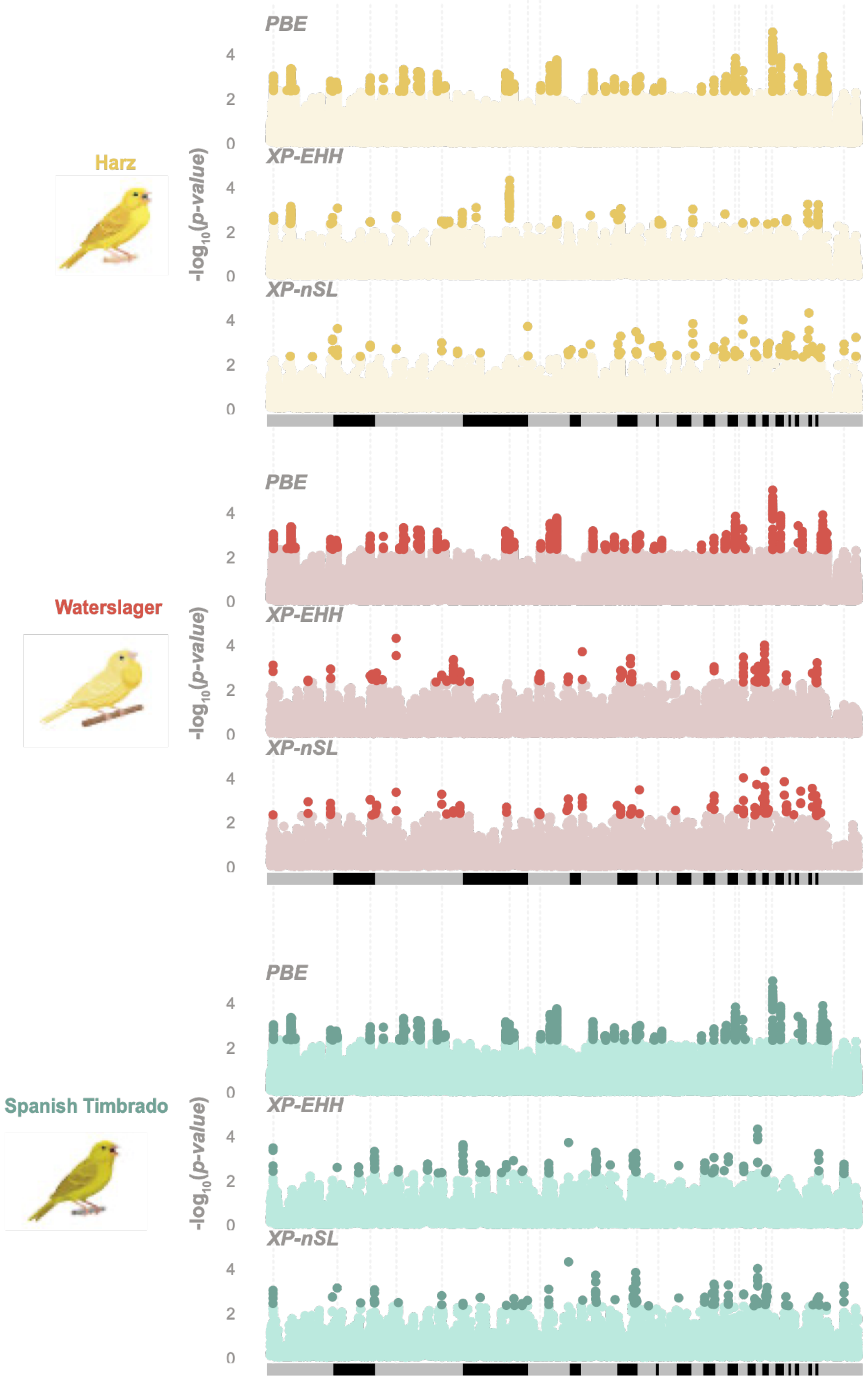


**Figure S4. Genome-wide signatures of selection across song-selected canary breeds.** The Manhattan plots represent each statistic used for DCMS, grouped by breed. Each point represents a 50 kb window, with the genomic position on the *x*-axis (chromosomes ordered according to the GCF_022539315.1 reference genome, with the exception that the unplaced scaffold of chromosome 7 was placed after chromosome 7). Points shown in the darker shade indicate windows falling within the top 0.5% of the distribution for each statistic. On the *y*-axis, the scores of each statistic were ranked and converted to *P*-values (see **Methods**), and the corresponding $-\log_{10}(P\text{-value})$ is shown. To facilitate comparison between the statistics and DCMS, the same annotated regions in **Fig. 3A** are highlighted here with vertical dashed lines.
